# *Burkholderia cenocepacia* and *Pseudomonas aeruginosa* coinfection increases host mortality by altering infection dynamics and compromising host immune effectiveness

**DOI:** 10.1101/2025.06.20.660666

**Authors:** Júlia Alcàcer-Almansa, Joana Admella, Núria Blanco-Cabra, Eduard Torrents

**Affiliations:** Bacterial infections and antimicrobial therapies group, Institute for Bioengineering of Catalonia (IBEC), The Barcelona Institute of Science and Technology (BIST), Baldiri Reixac 15-21, 08028 Barcelona, Spain; Microbiology Section, Department of Genetics, Microbiology and Statistics, Faculty of Biology, University of Barcelona (UB), 643 Diagonal Ave., 08028, Barcelona, Spain; Microbiology Section, Department of Biology, Healthcare and Environment, Faculty of Pharmacy and Food Sciences, University of Barcelona (UB), Av Joan XXIII, 27-31, 08028 Barcelona, Spain

**Keywords:** Coinfection, polymicrobial infection, *Galleria mellonella*, innate immunity, antibiotic susceptibility, virulence

## Abstract

Polymicrobial infections promote the appearance of a network of interactions that can lead to an increase in their antimicrobial tolerance or to the evasion of the host immune system. *Pseudomonas aeruginosa* and *Burkholderia cenocepacia* are two multidrug-resistant opportunistic pathogens that significantly influence host health and alter antibiotic responses when coinfected. To characterize host-pathogen dynamics, we examined infection progression, immune responses, bacterial virulence gene expression, global transcriptomic analysis, and antibiotic susceptibility in single and coinfections involving acute and chronic *P. aeruginosa* strains combined with *B. cenocepacia*. This work was performed *in vivo* using *G. mellonella* larvae as a model. Larval survival and bacterial dissemination were monitored, revealing tissue-specific patterns of infection. Our findings indicated that coinfections increased larval lethality and worsened overall host health. Notably, *B. cenocepacia* suppressed host melanization and immune responses, while *P. aeruginosa* triggered a strong immune activation. Coinfection also induced upregulation of virulence genes in both pathogens. This study advances understanding of host-pathogen interactions in polymicrobial infections and highlights the need for improved therapeutic strategies.

## Introduction

Microbial studies have traditionally focused on single-species populations. However, interspecies interactions within bacterial communities influence population dynamics, virulence, and responses to antimicrobials and the host immune system^1^. This is particularly relevant in cystic fibrosis (CF), an autosomal recessive disease caused by over 2,000 mutations in the cystic fibrosis transmembrane conductance regulator (CFTR) gene. The CFTR gene encodes an ion transporter channel, and its malfunction leads to a thickening of the extracellular mucus, thus preventing adequate mucociliary clearance^2^. This becomes an ideal environment for bacterial colonization by pathogens such as *Pseudomonas aeruginosa* and members of the *Burkholderia cepacia* complex (Bcc), which are associated with a high risk of pulmonary damage and often lead to chronic infection^3^. CF infections are polymicrobial, enabling interspecies interactions that can alter bacterial traits such as virulence and antimicrobial resistance while also modifying host-pathogen dynamics to evade the immune system^4–6^.

This study focuses on *P. aeruginosa* and *Burkholderia cenocepacia*, two major CF-associated pathogens. *P. aeruginosa*, an opportunistic Gram-negative pathogen, is prevalent in individuals with CF due to its adaptability and pathogenesis^7^. In this context, acute strains trigger rapid growth and inflammation, causing lung damage, whereas chronic strains persist, evade immunity, and induce progressive inflammation and lung function decline^8; 9^. On the other hand, *B. cenocepacia*, a Gram-negative pathogen, causes a severe reduction in lung function and potentially fatal systemic infections like cepacian syndrome. It is highly antibiotic-resistant and adapts to host immune responses and nutrient availability^10^. Despite their clinical importance, studies on interactions between these species remain limited, emphasizing the need for further research to better understand and address their coinfections.

*Galleria mellonella* was chosen as the *in vivo* model for this study of bacterial interactions due to its suitability for studying virulence and pathogenicity^11; 12^. *G. mellonella* presents several advantages such as its easy manipulation, its lack of nociceptors that lead to an absence of ethical regulations, and its simple anatomical structures that facilitate its infection, dissection, and infection monitoring^13^. A key feature of *G. mellonella* is its innate immune response, which shares similarities with that of mammals, consisting of a cellular and a humoral response^14^. Upon pathogen invasion, *G. mellonella* recognizes pathogen and damage-associated patterns through its pattern recognition receptors, which lead to the induction of signaling cascades such as the IMD pathway, mainly activated by Gram-negative bacteria^15; 16^. Then, the two components of the innate immune response, cellular and humoral, are activated. Cellular immunity is mediated by hemocytes, which are equivalent to neutrophils in mammals. They are predominantly found in the hemolymph, an analog of the bloodstream, and are implicated in pathogen phagocytosis, nodulation, and encapsulation. Besides, the humoral immune response includes the production of antimicrobial peptides (AMPs), lytic enzymes, melanization and oxidative stress. These processes are interconnected and highly relevant for microbial containment and infection clearance^17^.

This study aims to unravel the dynamics of *P. aeruginosa* and *B. cenocepacia* single- *vs.* coinfection in *G. mellonella,* an animal model of infection, focusing on bacterial virulence, host immunity, and infection dissemination. This work sheds light on understanding the differences between single and coinfection, as well as the roles of acute and chronic strains in this context, which brings us closer to a more adequate clinical response to polymicrobial infections.

## Materials and methods

### Bacterial strains and culture conditions

Strains used in this work are listed in Table S1. All strains were grown in Tryptic Soy Broth (TSB) medium (Scharlab).

### Bacterial strain preparation for G. mellonella infection

Strains were cultured overnight (ON) at 37 °C with agitation at 200 rpm. The ON bacterial suspension was centrifuged at 2500 *g* for 10 min (Labnet Spectrafuge™ 6C). Cell pellets were washed with 1X Phosphate-buffered saline (PBS) (Fisher Scientific) three times. The optical density at 590 nm (OD_590_) was measured and equalized to a final OD_590_ of 1 (10^8^ CFU) for *P. aeruginosa* acute and chronic strains, and 1.5 (10^6^ CFU) for *B. cenocepacia*, and subsequent dilutions were made to reach an inoculum concentration specified in each figure.

### G. mellonella maintenance, infection and health index monitoring

The protocol used was previously described^11^. Groups of 10 larvae (200-300 mg) were injected with different dilutions through the top-left proleg using a 26-gauge microsyringe (Hamilton), and injection with 1X PBS was used as a negative control. Larvae were monitored at 37 °C during the infection course. The different inoculum doses were plated on tryptic soy agar (TSA) (Scharlab) plates to determine bacterial CFU. For the antibiotic treatment survival curves, infected groups were treated with 40 mg per kg of larvae of ciprofloxacin (CPX) 1 h after infection. Larvae were monitored at 37 °C.

Infection evolution was monitored at two stages, labeled as early and late infection. Based on the larval survival shown in Figure S1 and previous work^12^, for *P. aeruginosa* PAO1 and coinfected groups, infection stages correspond to 5 h and 15 h for early and late infection stages, respectively. For the *B. cenocepacia* and *P. aeruginosa* PAET1 groups, the infection stages correspond to 15 h and 30 h for early and late infection, respectively. The health state of larvae was monitored visually and by gently touching them with a pipette tip, following the criteria described in Table S3. Health index scores were calculated as percentages of the maximum score.

### G. mellonella hemocyte and bacterial quantification

At the indicated time points, infected larvae were anesthetized on ice, hemolymph from 10 larvae per group was collected into Eppendorf tubes, and the hemolymph was kept on ice throughout the process. For bacterial quantification, serial dilutions of hemolymph were prepared, and CFU were determined by plating on TSA. To determine the number of hemocytes per mL of hemolymph, 80 μL of hemolymph per group were collected and centrifuged at 250 g for 5 min at 4 °C. Then, the supernatant was discarded, and the pellet was resuspended with 100 μL of 1X PBS and washed twice. Last, the pellet was resuspended in 50 μL of 1X PBS, and 10 μL of the suspension were mixed with 10 μL of Trypan Blue (Sigma-Aldrich). The cell concentration was determined by counting viable cells in a hematocytometer (BLAUBRAND) using an inverted fluorescence microscope (ECLIPSE Ti − S/L100, Nikon).

For hemocyte imaging, the washed hemolymph was stained with 5 μg/mL of FM^TM^ 4-64 (Invitrogen) and observed under an LSM 800 confocal laser scanning microscope (Zeiss). Images were analyzed using ZEN Microscopy Software (version 2.6) and ImageJ FIJI (version 1.52p).

### Phenoloxidase activity assay

At the indicated time points, the hemolymph of 10 larvae per group was extracted as described in the previous section. The hemolymph pool was centrifuged at 4000 g for 10 min at 4 °C. The supernatant was then collected and diluted at 1:3 (v/v) in 1X PBS at pH 6.5. In a 96-well plate (Costar 96-Well Black Polystyrene plate, Corning), 165 μL of 1X PBS at pH 6.5 and 10 μL of diluted hemolymph were added per well, and six replicates per condition were made. Then, 25 μL of a 20 mM solution of 4-methyl catechol (Sigma-Aldrich) were added to each well to start the enzymatic reaction. The absorbance of each well was measured at 490 nm every minute for 1 h in an Infinite 200 Pro Fluorescence Microplate Reader (Tecan) at room temperature.

### G. mellonella dissection

Infected larvae at the indicated time points were frozen at -20 °C for at least 24 h. Then, under a stereoscopic microscope (VWR) with the VisiLight CL-150 (VWR), larvae were cut with a sterile surgical blade, and a section of their gut and fat body was placed on top of microscope coverslips (Epredia 1,5 coverslips). Then, tissue samples were stained with FM^TM^ 4-64 (Invitrogen) at 5 μg/mL. Whole *G. mellonella* larvae images after dissection were taken with a mobile phone camera (Oppo A74) through the stereoscopic microscope (VWR). Stained tissues were observed under an LSM 800 confocal laser scanning microscope (Zeiss), and images were analyzed using ImageJ.

### G. mellonella whole larvae bacterial counts and total RNA extraction

At the indicated time points, 5 infected larvae per group were anesthetized on ice for 10 min in a 10 mL tube. Then, the larvae were disrupted and homogenized with a previously sterilized T 10 basic ULTRA TURRAX (IKA-Werke GmbH & Co.) homogenizer with an 8 mm diameter probe. To determine the bacterial CFU on each group, 100 mg of the mixture were plated in TSA. For samples with a high bacterial load, serial dilutions were prepared from 100 mg of the homogenized sample and plated on TSA to determine the number of CFU/mg of larvae. To extract RNA from the larvae and the bacteria, 140 mg of homogenized tissues at late infection timepoints were used for RNA extraction following the GeneJET RNA Purification Kit instructions (Thermo Fisher Scientific). The obtained RNA was treated with 10X TURBO DNase (Life Technologies) for 1 h to eliminate possible DNA contamination. The absence of DNA was verified by PCR amplification of the 18S ribosomal RNA (18S), *gyrase B* (*gyrB*), or *glyceraldehyde-3-phosphate dehydrogenase* (*gap*) housekeeping genes, using genomic DNA from *G. mellonella*, *B. cenocepacia,* or *P. aeruginosa*, respectively, as positive controls. RNA was quantified in a NanoDropTM 1000 spectrophotometer (Fisher Scientific).

### Reverse transcription and Real-time quantitative PCR

RNA to cDNA reverse transcription was performed using Maxima Reverse Transcriptase (Thermo Fisher Scientific) and Oligo (dt)18 primers or random hexamers (Thermo Fisher Scientific) according to manufacturer’s instructions. Quantitative real-time PCR (RT-qPCR) was performed using PowerUp^TM^ SYBR^TM^ Green Master Mix (Applied Biosystems) in a StepOnePlus^TM^ Real-Time PCR System (Applied Biosystems) using the primers stated in Table S2. The 18S, *gyrB* and *gap* genes were used as internal standards as their expression is vital and constant in *G. mellonella, B. cenocepacia* and *P. aeruginosa*, respectively. For each sample, three replicates were performed. The results were analyzed using the comparative Ct (cycle threshold) method (ΔΔCt). Previously published data was used to support the findings in this study, particularly on the *P. aeruginosa* and *B. cenocepacia* virulence genes in single- *vs.* coinfection ^18^.

### RNA sequencing library preparation

RNA was extracted from samples as described in the previous sections. Then, samples were sent to the NovoGene company (Munich, Germany) for further processing, RNA sequencing, and analysis. Briefly, ribosomal RNA (rRNA) was removed from total RNA using the Illumina Ribo-Zero Plus rRNA Depletion Kit prior to library preparation. RNA was subsequently fragmented and reverse-transcribed into cDNA using random hexamer primers, followed by second-strand synthesis. Strand-specific libraries were generated using dUTP incorporation and USER enzyme digestion. Libraries were prepared through end repair, A-tailing, adapter ligation, size selection, PCR amplification, and purification. Library quality was assessed using Qubit fluorometry, real-time PCR, and Bioanalyzer analysis.

Sequencing was performed on an Illumina NovaSeq X Plus platform in paired-end 150 bp mode (PE150), generating approximately 15 Gb of raw data per sample. Raw reads were processed using fastp to remove adapter sequences and low-quality bases. High-quality reads were aligned to bacterial reference genomes using Bowtie2, and to eukaryotic genomes using HISAT2. Reference genomes included *P. aeruginosa* PAO1 (NCBI RefSeq assembly GCF_000006765.1), *B. cenocepacia* J2315 (NCBI RefSeq assembly GCF_000009485.1), and *G. mellonella* (NCBI RefSeq assembly GCF_026898425.1). Gene-level read counts were obtained and used for downstream differential expression analysis. Differential expression was performed using DESeq2 (v1.42.0), with genes considered significantly differentially expressed at adjusted p-value ≤ 0.05 and |log2 fold change| ≥ 1. Functional enrichment analysis was performed using clusterProfiler. KEGG pathway enrichment analysis was performed on significantly differentially expressed genes. It is worth mentioning that conducting the differential expression analysis with the adjusted *p*-value resulted in fewer than 50 significant genes in the *P. aeruginosa* analysis, which did not allow for the KEGG enrichment analysis. For this reason, and as an exploratory approach, the non-adjusted *p*-value ≤ 0.05 was used to determine significance for *P. aeruginosa* PAO1.

### Data visualization and statistical analysis

All graphs, data analysis and statistical significance were made using Prism 10.1.1 (GraphPad Software).

## Results

### Coinfection reduces G. mellonella survival compared to single infections, with a minimal effect on antibiotic efficacy

*P. aeruginosa* and *B. cenocepacia*, both multidrug-resistant pathogens, are frequently associated with lung infections in CF patients. Previous studies using *G. mellonella* have shown altered survival outcomes in multispecies infections, although few studies have examined interactions between *P. aeruginosa* and *B. cenocepacia*^18; 19^. To identify suitable doses for coinfection experiments with *P. aeruginosa* and *B. cenocepacia*, larvae were first infected with different bacterial concentrations in single and coinfection (Figure S1). Then, to select the best representative strains, different acute (PAO1, PA54 and PA166 (Figure 1A)), and chronic (PAET1, PAET2 and PAET4 (Figure 1B)) *P. aeruginosa* strains (Table S1) were tested.

**Figure 1.**
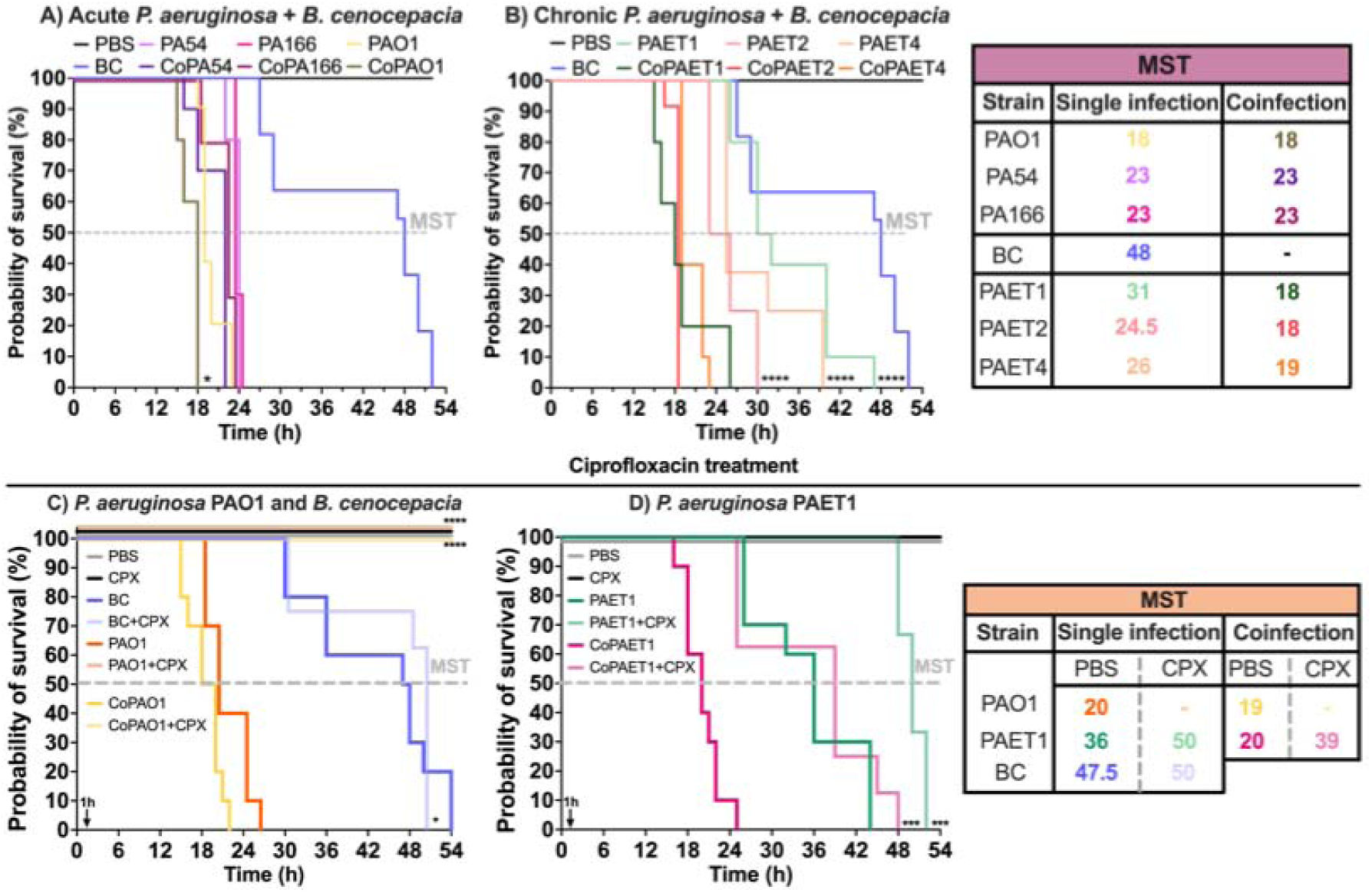
Survival curves of *G. mellonella* under coinfection and antibiotic treatment. **A)** Larval survival when infected with *B. cenocepacia*, acute *P. aeruginosa* strains, and their coinfection. **B)** Larval survival when infected with *B. cenocepacia*, chronic *P. aeruginosa* strains, and their coinfection. **C)** Ciprofloxacin (CPX) treatment of infected larvae with *P. aeruginosa* PAO1, *B. cenocepacia* and their coinfection. **D)** CPX treatment of infected larvae with *P. aeruginosa* PAET1 and its coinfection with *B. cenocepacia.* CPX was administered at a concentration of 40 mg/kg larvae 1 h after infection (see arrow). The following abbreviations were used: *B. cenocepacia* (BC), *P. aeruginosa* PA54 (PA54), *P. aeruginosa* PA166 (PA166), *P. aeruginosa* PAO1 (PAO1), *P. aeruginosa* PAET1 (PAET1), *P. aeruginosa* PAET2 (PAET2), *P. aeruginosa* PAET4 (PAET4), median survival time (MST, indicated with a dashed line in survival curves). Coinfection of *P. aeruginosa* - *B. cenocepacia* was noted as “Co” before the corresponding *P. aeruginosa* strain. Larvae were infected with defined doses (CFU/larva) of each strain: <10 for acute strains, 1x10^3^ for chronic strains, and 1x10^6^ for *B. cenocepacia*. Statistical significance was assessed using a log-rank (Mantel-Cox) survival test comparing each infected group with the coinfected group, with \**p*<0.05, \*\**p*<0.01, \*\*\**p*<0.001, and \*\*\*\**p*<0.0001.

Using Kaplan-Meier survival curves (Figure 1), we observed decreased larval survival following coinfection compared with single-infection groups. This effect was evident in larvae coinfected with chronic *P. aeruginosa* strains, with larger differences in median survival times (MST, the time at which 50% of individuals are no longer viable) between single and coinfected larvae. The MST differences in acute strains were less pronounced than in chronic strains. Given that the *P. aeruginosa* PAET1 strain exhibited the largest survival difference between single and coinfected groups (Figure 1B, 31 h *vs.* 18 h), it was selected as the representative chronic strain for this study. In contrast, larvae infected with acute *P. aeruginosa* strains exhibited a rapid loss of viability in both single- and coinfection conditions, with no MST differences between those groups. As a result, the effects of coinfection were likely masked in larvae infected with the acute *P. aeruginosa* PA54 and PA166 strains, because their short lifespans did not allow significant survival differences to emerge. However, although the MST was the same, significant differences were observed between single- and coinfected groups when using the *P. aeruginosa* PAO1 strain, which was therefore chosen as the representative acute strain for this study.

Larvae health state after infection was assessed using a scoring system (Table S3) based on activity, silk cocoon formation, melanization, and survival^20^. Assessments were performed at early (5 h for PAO1 and coinfected groups; 15 h for PAET1 and *B. cenocepacia*) and late (15 h for PAO1 and coinfected groups; 30 h for PAET1 and *B. cenocepacia*) post-infection stages. Larval health generally declined following infection, with the greatest impairment observed in coinfected larvae at early stages and in coinfected and PAO1-infected larvae at late stages compared with controls (Figure S2A). To correlate larval health status with their respective bacterial load, infected larvae were homogenized, and total bacterial loads quantified (Figure S2B). Interestingly, at late stages of infection, coinfected larvae exhibited lower *P. aeruginosa* loads in both acute and chronic strains compared to single infections (Figure S2B), while *B. cenocepacia* counts remained unchanged.

To test the impact of coinfection on antibiotic efficacy, larval viability following treatment with ciprofloxacin (CPX), administered 1 h post-infection with single or coinfection, was also evaluated. In the case of *B. cenocepacia*, CPX had minimal impact on larval survival (Figure 1C, blue lines). In single infections with the *P. aeruginosa* PAET1 chronic strain, CPX significantly delayed larval death, increasing MST from 36 h to 50 h (CPX) (Figure 1D). Even more pronounced effects were observed with *P. aeruginosa* PAO1, in which all antibiotic-treated larvae survived, in contrast to the untreated group, which had an MST of 20 h (Figure 1C, orange lines). Similar outcomes were observed in coinfections with *P. aeruginosa* PAO1, where CPX treatment resulted in 100% larval survival.

### Coinfection decreases melanization, promotes systemic dissemination, and inhibits the cellular immune response

Bacterial dissemination during infection was examined in the fat body, gut, and hemolymph using confocal microscopy (Figure 2). Early into the infection, *B. cenocepacia* and *P. aeruginosa* PAET1 (white and yellow arrows, respectively) were detected in both tissues during single infections and coinfections (also quantified in S4 Table), while *P. aeruginosa* PAO1 was undetectable, likely due to low bacterial load. At later infection stages (Figure 2, F-H rows), both pathogens were present in all tissues, with *P. aeruginosa* PAO1 (yellow arrows) showing widespread dissemination across the fat body and gut in both single and coinfected groups (Table S4). On the other hand, bacterial quantification in each tissue allowed us to determine that *B. cenocepacia* tended to accumulate in the gut (Figure 2, 1C and 1G and Table S4), while both strains of *P. aeruginosa* preferentially accumulated in the fat body (Figure 2, 2-3B and 2-3G, and Table S4).

**Figure 2.**
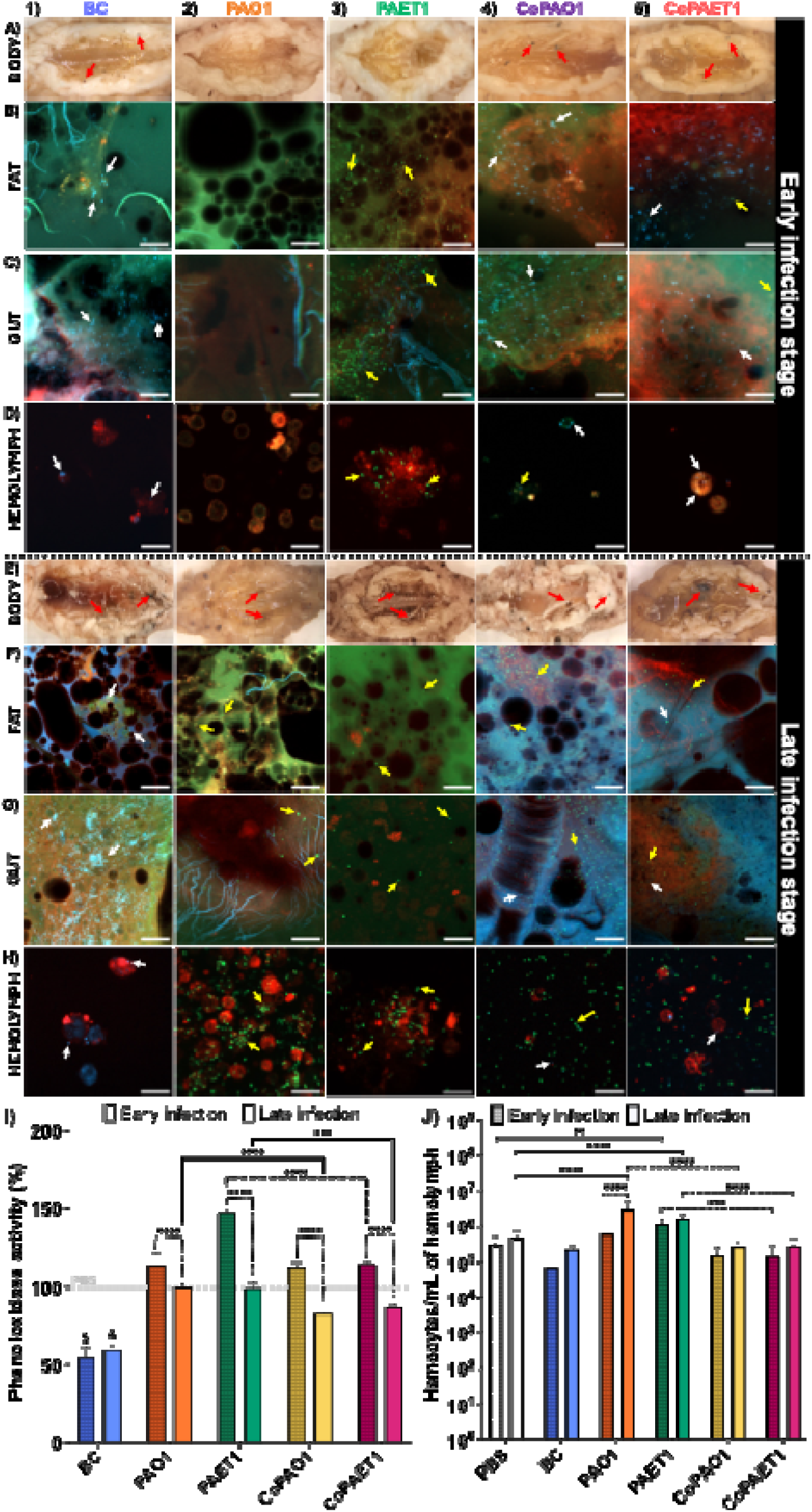
*G. mellonella* melanization, bacterial dissemination and cellular immune response in infected larvae. **1)** *B. cenocepacia* (BC) **2)** *P. aeruginosa* PAO1 (PAO1) **3)** *P. aeruginosa* PAET1 (PAET1) **4)** Coinfected groups with PAO1 and BC (CoPAO1) **5)** Coinfected groups with PAET1 and BC (CoPAET1). **A)** and **E)** show images of dissected larvae under the stereoscopic microscope at early and late infection time points (BODY). Melanization can be noted as black spots in dissected larvae, highlighted with red arrows. Images of larval fat body (**B** and **F)** and gut (**C** and **G**) were taken under the confocal microscope at early and late infection time points, respectively. **D)** and **H)** show confocal images of the hemolymph from the different groups at early and late times of infection. In the confocal images, PAO1 and PAET1 can be seen in green and highlighted with yellow arrows, BC can be seen in blue and highlighted with white arrows, and hemocytes are seen in red. Scale bar corresponds to 20 μm. **I)** corresponds to the hemolymph phenoloxidase activity assay of different infection conditions compared to the control. Data is presented as the mean ± SD for each condition. The control baseline (PBS) is indicated with a gray dashed line. **J)** shows the number of hemocytes (cells/mL) in the hemolymph. Data is presented as the mean ± SD for each condition. Statistical significances were evaluated and established through a two-way ANOVA, where \**p*<0.05, \*\**p*<0.01, \*\*\**p*<0.001 and \*\*\*\**p*<0.0001. Significant differences between individual groups *vs.* coinfection at early stages are shown as a dashed line, and at late stages as a continuous line. $ and & correspond to significant differences (****) in all comparisons between BC and the rest of the groups at early and late infection stages, respectively.

Melanization is a critical component of the *G. mellonella* immune response, triggered by microbial recognition and resulting in melanin deposition that encapsulates pathogens and limits the spread of infection^21^. The melanization response was first monitored by larval dissection and observation under a stereoscopic microscope (Figures 2A and E), in which *P. aeruginosa* induced a more generalized melanization, while *B. cenocepacia* infections produced localized patterns at late stages of infection (Figure 2E). Coinfected larvae display a combined melanization phenotype (Figures 2, 4-5E), with larger melanization spots and widespread body blackening corresponding to reduced general body melanization. To dig deeper into the larval melanization response, melanization capacity after infection was evaluated by monitoring phenoloxidase activity ^22^ in hemolymph samples at early and late infection stages (Figure 2I). Even though some melanization spots were observed, phenoloxidase activity was impaired during infections with *B. cenocepacia*, particularly in single infections (Figure 2I, blue bars). This can be observed by the appearance of larvae immediately after death, in which *B. cenocepacia*-infected larvae retained a coloration similar to that of living individuals (Figures 2 1E and S3 C). In contrast, larvae infected with *P. aeruginosa* showed enhanced phenoloxidase activity (Figure 2I, green striped bar and Figure 2 3E).

Furthermore, to explore the *G. mellonella* cellular immune response, hemocyte concentration in hemolymph samples was determined (Figure 2J). Single *P. aeruginosa* infections significantly increased hemocyte counts compared to the control, with the acute *P. aeruginosa* PAO1 strain (Figure 2J, orange bars) inducing the highest hemocyte proliferation. When focusing on bacterial growth, *P. aeruginosa* PAO1 highly proliferates in the hemolymph (Table S3) and does not appear to be phagocyted (Figure 2 2H, 4-5H). In contrast, this study also reveals a decrease in hemocyte concentration when *B. cenocepacia* was present in single infections (Figure 2J, blue bars). Remarkably, hemocyte proliferation during coinfection resembled that observed with *B. cenocepacia* alone (Figure 2J, yellow and magenta bars). When observing the hemolymph under the confocal microscope, *B. cenocepacia* was found exclusively inside the hemocytes (Figure 2 D and H, white arrows). Also, although there are no significant differences in hemocyte concentration between larvae infected with *B. cenocepacia* and the control (Figure 2J), the hemocytes were morphologically altered, with increased internal compartmentation and increased size. This can also be seen by comparing the infected hemocytes of Figure 2D and 2H with the non-infected group (Figure S3B).

### Differential immune gene activation reflects pathogen-dependent immune modulation

To further investigate larval immune responses, the expression of various immunity-relevant genes in *G. mellonella* was analyzed through RT-qPCR. The expression of the gene encoding for the transcription factor Relish, involved in pathogen recognition, was predominantly repressed at early stages, becoming overexpressed at later infection stages across all *P. aeruginosa*-infected groups (Figure 3A, orange, green, yellow and magenta solid bars). However, *B. cenocepacia* in single infection repressed the expression of this gene at both infection stages (Figure 3A, blue striped and solid bars). A similar pattern was observed when studying the *nos* gene, encoding nitric oxide synthase (NOS), which is responsible for creating oxidative stress to control the infection (Figure 3B). Notably, *P. aeruginosa* PAO1 induced the highest expression levels for the genes encoding for Relish and NOS in both single and coinfected larvae.

**Figure 3.**
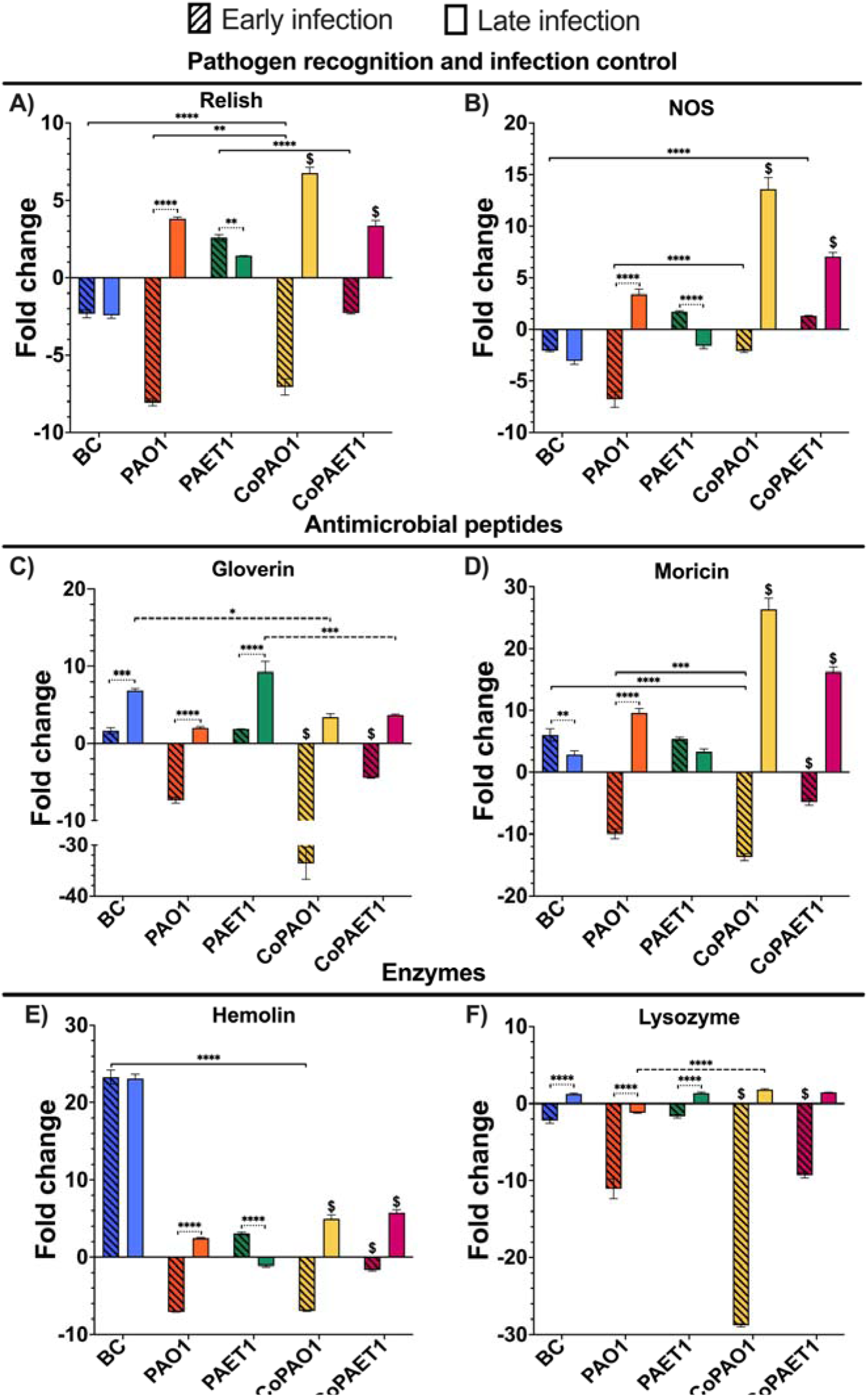
Expression of *G. mellonella* immune-relevant genes under different infection conditions. The fold change expression is plotted in relation to the control (larvae injected with PBS). Data is presented as the mean ± SD of the fold changes for each condition. The following abbreviations were used: *P. aeruginosa* PAO1 (PAO1), *P. aeruginosa* PAET1 (PAET1), *B. cenocepacia* (BC). Coinfection was noted as “Co-” before the corresponding strain. For early and late infection timepoints, see materials and methods. Statistical significance was established through a two-way ANOVA. $ corresponds to significant differences (****) in all comparisons with other groups, where \**p*<0.05, \*\**p*<0.01, \*\*\**p*<0.001 and \*\*\*\**p*<0.0001.

The genes encoding for the AMPs Gloverin and Moricin appeared over-expressed in all conditions at late infection stages (Figure 3C-D). However, at early stages, they were repressed in larvae infected with *P. aeruginosa* PAO1 (Figure 3C-D, orange striped bars) and in coinfected larvae (yellow and magenta striped bars). For Gloverin, gene expression levels stand out for *B. cenocepacia* and *P. aeruginosa* PAET1 (Figure 3C, blue and green bars). In contrast, their coinfection resulted in a lower induction, comparable to the levels seen with *P. aeruginosa* PAO1 and its coinfection (Figure 3C-D, orange and yellow bars). Regarding Moricin, *B. cenocepacia* single infection reduces its gene expression over time, similar to that observed in *P. aeruginosa* PAET1-infected larvae. Again, the behavior of *P. aeruginosa* PAO1 and the coinfected groups was similar, inducing higher gene expression in their late stages of infection. In addition, CoPAO1 (Figure 3D, yellow solid bar) stands out as the condition with the highest Moricin gene induction (fold change of 26.3), whereas the expression levels of the genes encoding for Gloverin and Moricin in the *B. cenocepacia*-infected groups were notably lower. Last, the gene encoding for the immunoglobulin-like protein Hemolin appeared highly upregulated in *B. cenocepacia* single infections, while its expression in coinfection was only upregulated at late infection stages for both *P. aeruginosa* strains.

To better understand the molecular mechanisms behind the increased larval mortality during coinfection, we conducted a comprehensive global transcriptomic analysis using RNA sequencing. This analysis was performed on larvae infected with *P. aeruginosa* PAO1, *B. cenocepacia*, or both pathogens (Figures 4 and 6). We examined changes in the transcriptomes of *G. mellonella, B. cenocepacia*, and *P. aeruginosa* PAO1. *P. aeruginosa* PAET1 was excluded from the analysis due to the lack of a reference genome, which prevented accurate read alignment and transcriptomic analysis.

**Figure 4.**
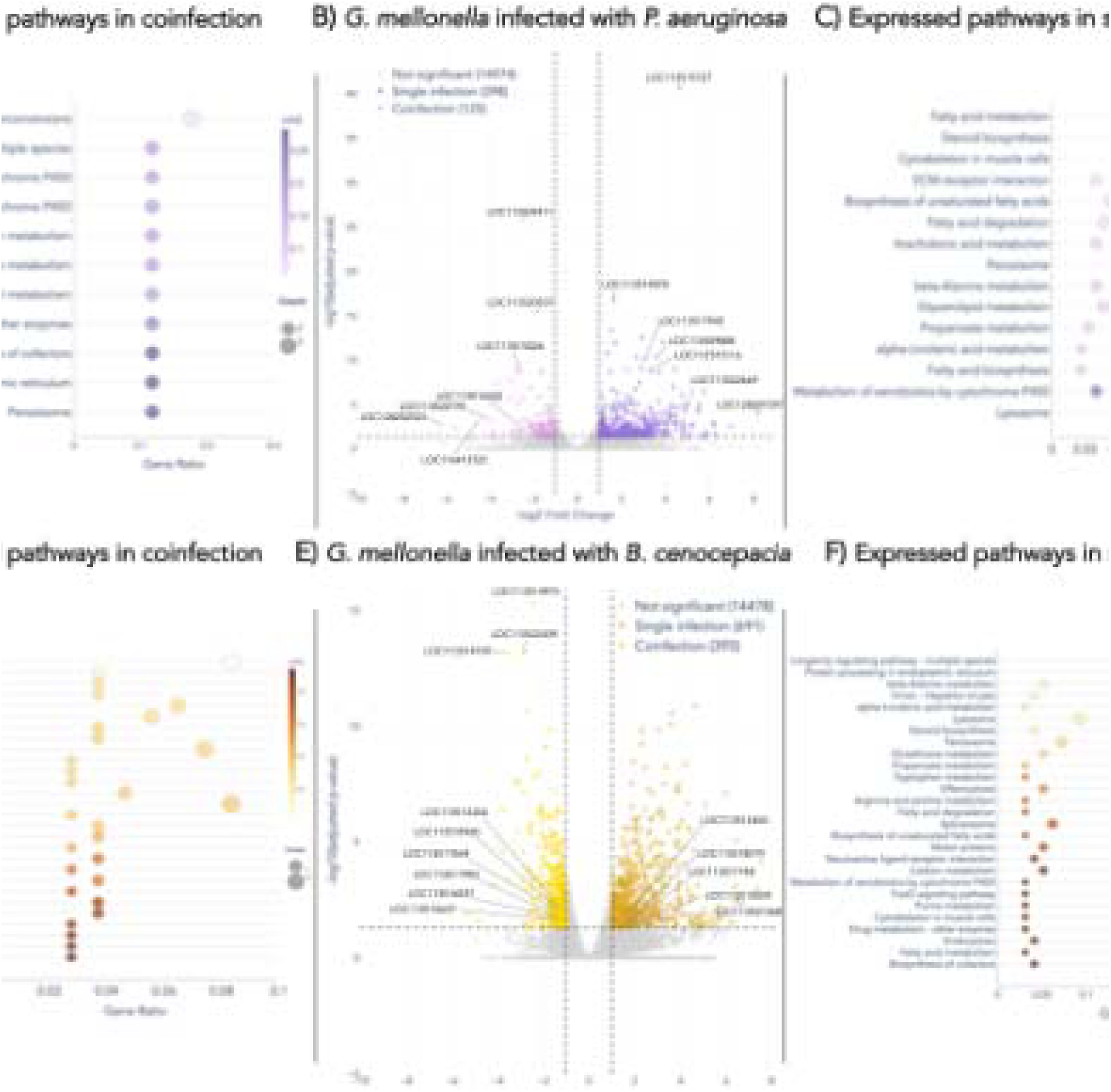
Differential gene expression of *G. mellonella* during single infection and coinfection with *P. aeruginosa* PAO1 or *B. cenocepacia*. Dot plots show the Kyoto Encyclopedia of Genes and Genomes (KEGG) pathway classification of the most enriched pathways for *G. mellonella* after infection with *P. aeruginosa* and *B. cenocepacia* in coinfection **(A** and **D)** and single infection **(C** and **F)**. The gene ratio is defined as the number of genes associated with each pathway divided by the total number of genes. The adjusted *p*-value for each pathway is represented by the color gradient, while the size of each dot indicates the number of genes in the corresponding pathway. **B)** and **E)** Volcano plots showing differentially expressed genes for *G. mellonella* after *P. aeruginosa* or *B. cenocepacia* coinfection (pink or light yellow) and single infection (purple or dark yellow), using a threshold of adjusted *p-*value < 0.05. Selected immune-related genes are annotated by locus tag.

**Figure 5.**
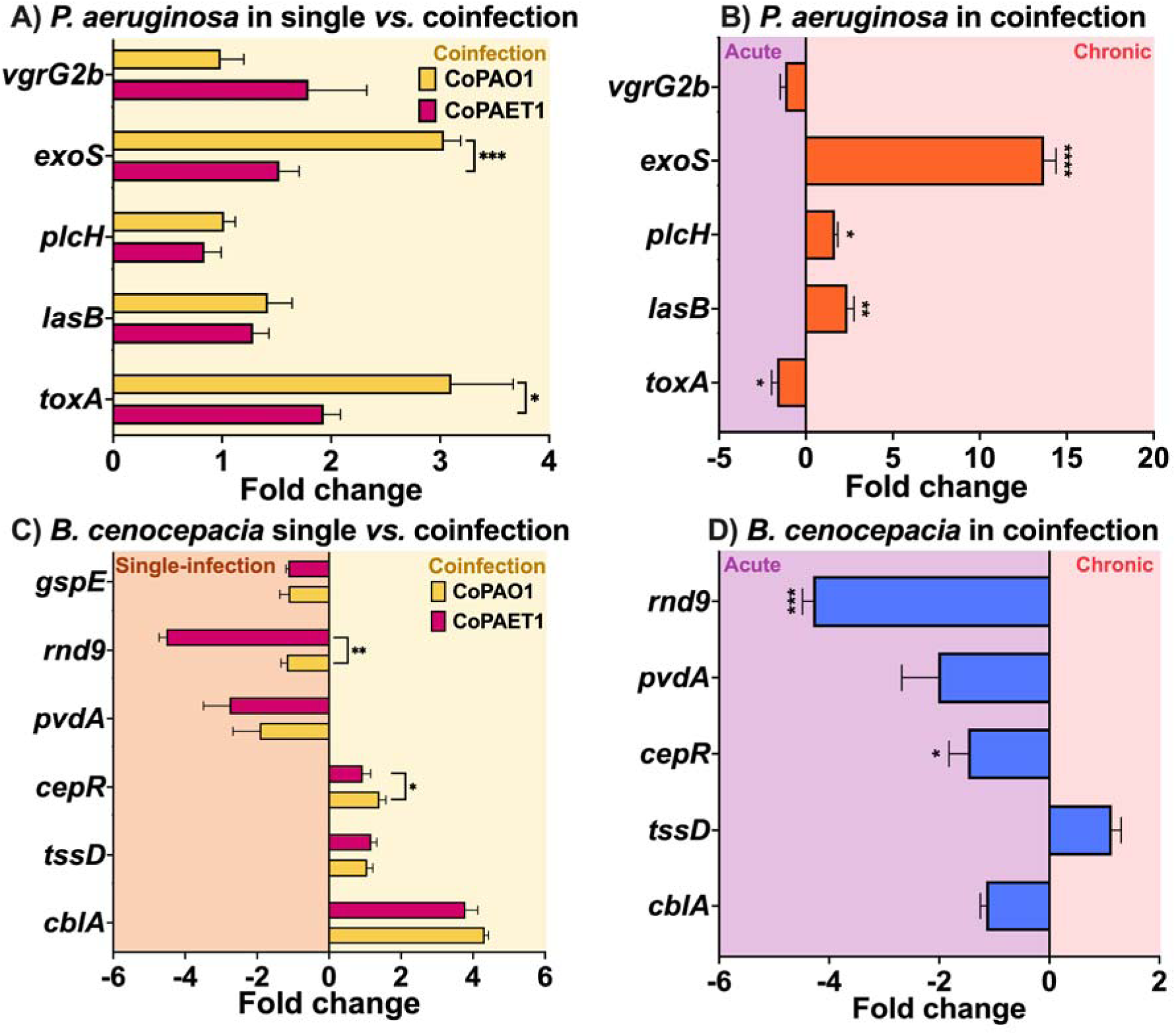
Bacterial virulence gene expression of single- *vs.* coinfection and acute *vs.* chronic strains in coinfection. **A)** *P. aeruginosa* virulence genes in single- and coinfection. Published data was used to support this study^18^. **B)** *P. aeruginosa* virulence genes of chronic *vs.* acute strains in coinfection. **C)** *B. cenocepacia* virulence genes in single- and coinfection. Published data was used to support this study^18^ **D)** *B. cenocepacia* virulence genes in coinfection with chronic *vs.* acute strains. Data is presented as the mean ± SD of the fold changes for each gene for chronic *vs.* acute strains in coinfection. Statistical significance was established through Student’s *t*-tests comparing the ΔCt of CoPAET1 vs. CoPAO1 for each gene, or the ΔΔCt of single- *vs.* coinfection, where \**p*<0.05, \*\**p*<0.01, \*\*\**p*<0.001 and \*\*\*\**p*<0.0001.

**Figure 6.**
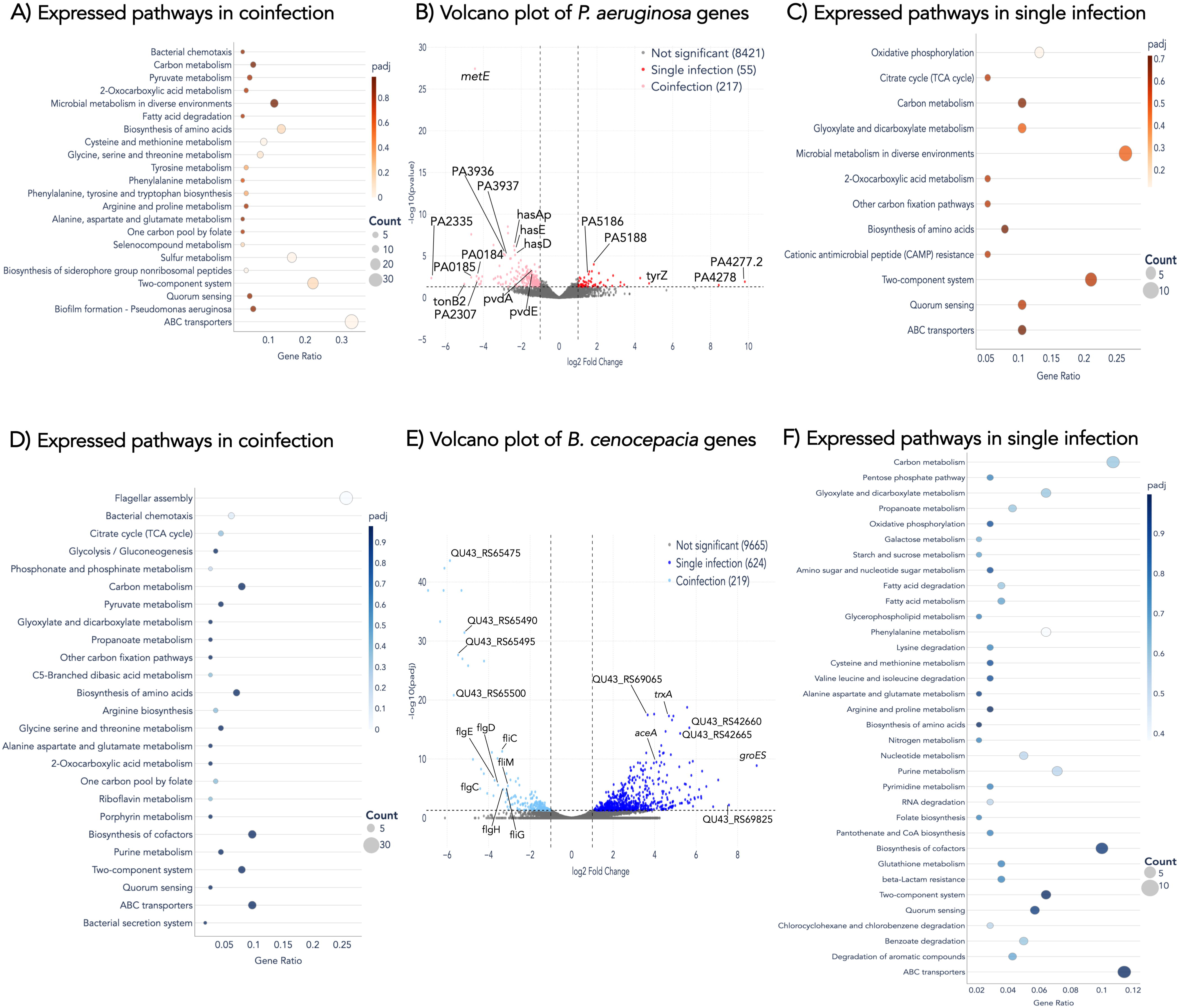
Differential gene expression of *P. aeruginosa* and *B. cenocepacia* in *G. mellonella* during single infection and coinfection with *P. aeruginosa* PAO1. Dot plots show the Kyoto encyclopedia of genes and genomes (KEGG) pathway classification of the most enriched pathways for *P. aeruginosa* and *B. cenocepacia* in coinfection **(A** and **D)** and single infection **(C** and **F)**. The gene ratio is defined as the number of genes associated with each pathway divided by the total number of genes. The adjusted *p*-value for each pathway is represented by the color gradient, while the size of each dot indicates the number of genes in the corresponding pathway. **B)** and **E)** Volcano plots showing differentially expressed genes for *P. aeruginosa* or *B. cenocepacia* coinfection (light red or light blue) and single infection (dark red or dark blue), using a threshold of adjusted *p-* value < 0.05. Selected virulence-related genes are annotated by gene name when available, or by locus tag when no functional annotation has been assigned.

By comparing the *G. mellonella* transcriptional profiles of larvae infected with *P. aeruginosa* PAO1 in single infection (Figure 4B, purple) versus coinfection with *B. cenocepacia* (Figure 4B, pink), a total of 523 differentially expressed genes (DEGs) were identified. Of these, 398 genes were overexpressed under single-infection conditions, indicating a pronounced shift in host gene expression in the absence of *B. cenocepacia*. KEGG pathway enrichment analysis revealed that DEGs specific to single infection were predominantly associated with fatty acid metabolism, steroid biosynthesis, and cytoskeletal organization (Figure 4C). Focusing on immunity-related genes in the comparison between *P. aeruginosa* PAO1 single infection and coinfected larvae (Table S5), several immune effectors and regulatory proteins were differentially expressed. In single infection, phenoloxidase-related genes (LOC113517945 and LOC113515116) were significantly modulated, along with LOC113515727, encoding a putative hemolectin, and multiple putative metalloproteases (LOC113521879, LOC113522639, LOC113522669). In contrast, coinfected larvae showed increased differential expression of canonical antimicrobial peptides, including gloverins (LOC113510422, LOC128201089), lysozyme (LOC113515290), moricin (LOC113509608), and gallerimycin (LOC113523440), consistent with patterns observed in Figure 3.

When comparing larvae infected with *B. cenocepacia* alone versus coinfection with *P. aeruginosa* PAO1, 1,084 DEGs were identified, with 691 genes differentially expressed under single-infection conditions (Figure 4E). KEGG pathway analysis indicated that the Toll and Imd signaling pathway was the most enriched pathway in coinfected larvae (Figure 4D), whereas longevity-regulating pathways were predominantly represented in single infection (Figure 4F). Among immune-related genes in this comparison, apomucin-like proteins (LOC113517748) and a putative gamma-interferon-inducible lysosomal thiol reductase-like protein (LOC113518019) were differentially expressed in single-infection conditions. In contrast, coinfected larvae exhibited differential expression of multiple components of the Toll pathway, including spaetzle genes (LOC113516037, LOC113513204, LOC113511546), as well as genes involved in oxidative stress responses, such as catalases (LOC113511903) and peroxidases (LOC113518960).

### Coinfection alters virulence gene expression in P. aeruginosa and B. cenocepacia

To assess differences in virulence gene expression between the reference strain *P. aeruginosa* PAO1 and the chronic isolate *P. aeruginosa* PAET1, RT-qPCR was employed to monitor the expression of selected virulence genes*. P. aeruginosa* and *B. cenocepacia* virulence-related genes (also at late infection stages), comparing single *vs*. coinfected larvae (Figure 5A and C), as well as acute (PAO1) *vs.* chronic (PAET1) *P. aeruginosa* strains in coinfected larvae (Figure 5B and D). Overall, both acute and chronic strains showed a tendency for increased virulence gene expression during coinfection, with *P. aeruginosa* PAO1 (the acute strain) exhibiting the most significant increase in virulence-related genes (Figure 5A). Further comparison of chronic versus acute strains in coinfection (Figure 5B) revealed that, interestingly, *lasB*, encoding the virulence-related enzyme Elastase B^23^, *plcH*, encoding Hemolytic phospholipase C^24^, and *exoS*, encoding the Type III Secretion System effector Exoenzyme S^25^, were more highly expressed in the chronic strain *P. aeruginosa* PAET1 compared to the acute strain. On the contrary, expression of *vgrG2b,* encoding Type VI secretion system spike protein VgrG2b^26^ and *toxA*, encoding Exotoxin A, were greater in the acute strain *P. aeruginosa* PAO1.

For *B. cenocepacia*, coinfection triggered upregulation of the virulence-associated genes *cepR,* encoding the CepR quorum-sensing receptor^27^, and *cblA*, whereas the antibiotic tolerance gene *rnd9*, encoding the RND9 efflux pump^28^, was more strongly expressed in single infections (Figure 5C). Within them, the expression of *cepR* and *rnd9* were statistically more expressed in acute infection rather than in chronic infection (Figure 5D), while *tssD*, another virulence-associated gene encoding a Type VI Secretion System subunit^29^, was more expressed in the chronic PAET1 coinfection.

To further characterize global transcriptional changes associated with single-species and coinfection conditions, RNA sequencing was conducted on late-stage infection samples from *G. mellonella* larvae infected with *P. aeruginosa* PAO1, *B. cenocepacia*, or both species. DEG analyses were performed by comparing single infections with coinfections.

By comparing the transcriptional profile of *P. aeruginosa* PAO1 in single infection (Figure 6B, light red) versus that of *P. aeruginosa* in coinfection (Figure 6B, dark red), 362 differentially expressed genes were found. The 217 genes overexpressed in coinfection (Figure 6B, light red) indicate a clear shift in gene expression when *B. cenocepacia* participates in infection. Interestingly, differentially expressed genes in coinfection were mostly associated with general metabolism, virulence, motility, and transport, according to KEGG categories (Figure 6A). Particularly, ABC transporters, including PA3936, PA3937, PA0184, PA0185, PA2307, *hasAp, hasE* and *hasD*, appeared as one of the largest and most relevant groups of genes overexpressed in *P. aeruginosa* PAO1 coinfection. Similarly, genes involved in quorum sensing, particularly *braD, lasI* and *rhll* (log2 fold change of -1.2, -1.5, and -1.5, respectively) and two-component system genes *pfeS* and *pfeR* (log2 fold change of -1.36 and -1.34, respectively) were altered when both bacterial species coexisted, indicating an increase in *P. aeruginosa* PAO1 virulence when in coinfection. Since the genes overexpressed in coinfection were extensive, a division was made based on their roles in virulence-related pathways, including secretion systems, siderophores, quorum sensing, virulence, and antibiotic resistance (Table S6). Within this classification, the numerous groups of upregulated siderophores and hemophores, including the entire pyoverdine gene family (*pvdA-S*), as well as genes involved in iron acquisition and transport, such as PA2335, *tonB2* and *pirA* (log2 fold change of -1.36), were highlighted. Among the 95 genes that were overexpressed in *P. aeruginosa* single infection, the genes with the highest log2 fold change correspond to PA4277.2 and PA4278, which encode a putative glycine transfer RNA (tRNA-Gly) and a putative TyrZ tyrosine–tRNA ligase, respectively. Among the most significant genes, PA5186 and PA5188 encode enzymes involved in fatty acid β-oxidation, including a putative acyl-CoA dehydrogenase and a putative 3-hydroxyacyl-CoA dehydrogenase, respectively, which potentially indicate that *P. aeruginosa* is metabolizing fat in *G. mellonella*. However, no direct correlation was found between their expression and virulence or host-pathogen interactions. It is also worth noting that most of the genes overexpressed in single infection are linked to metabolism, including the glyoxylate and oxidative phosphorylation pathways, with genes such as *acnA*^30^, which encodes aconitase (Figure 6C).

The differential gene profile was much broader for *B. cenocepacia*, with 845 differentially expressed genes between single-infection (Figure 6E, dark blue) and coinfection (Figure 6E, light blue). Unlike *P. aeruginosa*, a widely studied bacterial species, only a few genes have been characterized in *B. cenocepacia*. When focusing on the *B. cenocepacia* genes that were overexpressed in coinfection (Figure 6E, light blue), the described genes with a higher adjusted *p*-value were QU43_RS65475 (BCAM2221), encoding a putative RhtX/FptX family siderophore transporter^31^, QU43_RS65490 (BCAM2224), encoding a putative TonB-dependent siderophore receptor, and QU43_RS65495 (BCAM2225), encoding a putative ABC transporter ATP-binding protein^32^. The only significant pathways identified by KEGG analysis were flagellar assembly and bacterial chemotaxis, with 29 and 7 genes overexpressed, respectively, compared with single infections (Figure 6D). Among the characterized genes involved in flagellar assembly, the *fli* and *flg* families stand out and have been associated with virulence in various bacterial species, including *Burkholderia pseudomalle*^33^. On the other hand, *cheA* and *cheW*, encoding for a histidine kinase and an adapter protein, and with a log2 fold change of -2.45 and -2.6, respectively, were the only characterized genes in the bacterial chemotaxis classification. Interestingly, some characterized virulence-related genes were also overexpressed in coinfection. These include *cblA* (log2 fold change of -1.5), which encodes for the cable pilus major subunit of *B. cenocepacia*^34^, *eco* (log2 fold change of -1.4), which encodes for a serine protease inhibitor^35^, *tpx* (log2 fold change of -1.6), which encodes for a thiol peroxidase^36^, and *gspD* and *gspG* (log2 fold change of -1.3 and -1.27, respectively), encoding for the Type-II secretion system (T2SS) proteins GspD and GspG^37^. In addition, iron homeostasis genes were upregulated in coinfection. The *fhuB* gene (log2 fold change of -1.1), encoding for the Fe(3+)-hydroxamate ABC transporter permease FhuB^38^, as well as *hemD* and *hemF* (log2 fold change of - 1.13 and -1.23, respectively), encoding for a uroporphyrinogen-III synthase and a coproporphyrinogen-III oxidase, which participate in the heme biosynthesis pathway^39^, were also overexpressed in coinfection conditions.

The genes with the highest log2 fold change in single infection were *groES* and QU43_RS69825 (BCAS0229), encoding the co-chaperonine GroES and a putative ABC transporter substrate-binding protein, respectively (Figure 6B, dark blue). Other upregulated genes included *trxA* and *trxB* (log2 fold change of 2.4), encoding a thioredoxin and a thioredoxin-disulfide reductase, the latter of which has also been described to be overexpressed in virulence contexts^40^. When focusing on carbon metabolism-related pathways, including *aceA*, *pgl* (log2 fold change of 2.4), and *zwf* (log2 fold change of 1.8), which had been previously described to contribute to bacterial virulence in other species, were significantly overexpressed in *B. cenocepacia* single infection^41–44^. Additionally, phenylalanine metabolism appeared as one of the most relevant pathways in the KEGG gene classification (Figure 6F), revealing upregulation of the phenylacetic acid catabolic genes (*paaD*, *paaE*, *paaG*, *paaI*, *paaK*)^45^.

## Discussion

*P. aeruginosa* and *B. cenocepacia* are two major pathogens associated with CF that significantly compromise both the quality of life and life expectancy of affected individuals. In this study, we conducted a comprehensive analysis of interactions among these pathogens during coinfection, evaluated their combined impact on the *G. mellonella* host model, and compared *P. aeruginosa* acute and chronic strains.

Survival analysis revealed a statistically significant reduction in larval viability during coinfection compared with single infection, particularly with chronic *P. aeruginosa* strains. In contrast, acute *P. aeruginosa* strains caused rapid larval death in both single and coinfection conditions, with no difference in MST between the two (Figure 1). These findings are consistent with the known aggressiveness and rapid progression of strains isolated from acute infections, as both single and coinfected larvae with *P. aeruginosa* PAO1 showed the fastest mortality. In contrast, infections with the chronic clinical isolate *P. aeruginosa* PAET1, derived from a CF patient, showed a significantly lower mortality rate, highlighting the persistent characteristics of the chronic strain.

The presence of multiple bacterial species in coinfection can modulate antibiotic tolerance through interspecies interaction^47–49^. Previous *in vitro* studies have shown increased antibiotic sensitivity in dual-species biofilms of *P. aeruginosa* and *B. cenocepacia*^18^. Based on these findings, we aimed to evaluate whether this pattern is maintained in an *in vivo* model (Figure 1). It is important to note that the inoculum concentrations of *P. aeruginosa* PAO1 and *P. aeruginosa* PAET1 are 10 and 10³ CFU/larva, respectively, which may contribute to the higher survival rate observed in treated larvae infected with *P. aeruginosa* PAO1 compared to those infected with PAET1. However, because CPX is ineffective against *B. cenocepacia,* differences in inoculum size alone cannot account for the complete survival of larvae coinfected with *P. aeruginosa* PAO1 and *B. cenocepacia* after CPX treatment. This suggests that *P. aeruginosa* PAO1 is the primary driver of larval mortality in this coinfection, as seen in other works^19^.

Considering these initial findings on distinct larval survival, differences in the immune response between single-infected and coinfected larvae were assessed. AMPs Gloverin and Moricin, which are unique to Lepidoptera and serve as key components of the first line of defense against microbial infections^50^, were upregulated in larvae infected with *P. aeruginosa* PAO1 and in coinfected groups at later stages (Figure 5), consistent with previous findings involving *Mycobacterium abscessus* and the *P. aeruginosa* PAO1 and PAET1 strains^6^. These genes, as well as those encoding a putative Gallerimycin and Lysozyme, were also differentially expressed in coinfection, as revealed by RNA sequencing (Figure 4), corroborating these results. Interestingly, the chronic *P. aeruginosa* PAET1 strain triggered an early immune response across most of the virulence genes studied, whereas the acute *P. aeruginosa* PAO1 strain elicited a delayed but stronger response. These findings align with the existing literature, suggesting that chronic strains provoke an early immune response that eventually weakens as the pathogen adapts to the host^51^. This pattern may account for both the delayed mortality observed in *P. aeruginosa* PAET1 single-infected larvae (Figure 1) and the downregulation of immunity-related genes during the early stages of infection. However, given the relatively short duration of infection in *G. mellonella*, the extent to which host adaptation contributes to these responses should be interpreted with caution.

Melanization, a key component of the *G. mellonella* immune response to contain infection, was observed as a general blackening when larvae were infected with both *P. aeruginosa strains,* whereas *B. cenocepacia* infections typically resulted in localized melanization spots (Figure 2A and E). Assessment of melanization via phenoloxidase activity, a key enzyme in the pathway, revealed reduced enzymatic activity when *B. cenocepacia* was present during infection (Figure 2I). This pattern was also observed in RNA sequencing data, in which phenoloxidase transcripts were differentially expressed in larvae with a single infection of *P. aeruginosa* compared with coinfected larvae (Figure 4). These findings align with the ability of the *Burkholderia* genus to evade immune responses by producing surface lipopolysaccharides that provide physical protection against the immune system^52^. This ability may be linked to the downregulation of the gene encoding for Relish and NOS, key regulators of pathogen recognition and infection containment (Figure 3).

Hemocytes mediate the cellular immune response of *G. mellonella*, with their numbers fluctuating in response to pathogen exposure^11^. Their increased abundance during *P. aeruginosa* PAO1 infection (Figure 2J) likely reflects the strong immune activation triggered by the high bacterial loads observed in the hemolymph (Table S4). In contrast, *B. cenocepacia* infections resulted in reduced hemocyte levels (Figure 2J), possibly due to its ability to induce macrophage death, as described for other *Burkholderia* species^53; 54^. Notably, *B. cenocepacia* was consistently found inside the hemocytes (Figure 2 D and H), in agreement with studies demonstrating its intracellular survival in murine macrophages infected with *B. pseudomallei*^55^. Moreover, infected hemocytes displayed morphological alterations, suggesting cellular impairment associated with bacterial colonization. A possible explanation is the ability of *B. cenocepacia* to delay phagosomal maturation, weakening the innate immunity^56; 57^. Supporting this interpretation, expression of the gene encoding for Hemolin, an immunoglobulin-like opsonin involved in phagocytosis^58^, was upregulated (Figures 3 and 4). This is consistent with the intracellular nature of *B. cenocepacia* in the hemocytes observed in Figure 2, as its opsonin activity enhances phagocytosis of foreign cells^16; 58^. Additionally, a single infection of *B. cenocepacia* led to a downregulation of the gene encoding for Relish (Figure 3), a transcription factor that regulates the synthesis of antimicrobial peptides that combat Gram-negative pathogens^59; 60^, and a generally low activation of immune-related genes (Figure 3). These findings indicate that the presence of *B. cenocepacia* jeopardizes the host’s ability to overcome the infection.

Studying bacterial dissemination throughout the larvae provides valuable insights into infection progression and host-pathogen interactions. The fat body plays a crucial role in larval development and immune response, serving as an energy reserve, a site for lipid and glycogen synthesis, and AMP production^61^. Similarly, while the gut primarily handles digestion and absorption, it also functions as a key barrier contributing to immune defense^62^. While all pathogens were detected in both the fat body and the gut, *P. aeruginosa* was predominantly localized in the fat body, consistent with previously published findings^12^. This is also consistent with an upregulation of genes involved in β-oxidation and the glyoxylate shunt in *P. aeruginosa* during infection, as seen with the *acnA* gene differential expression in the transcriptomic analysis (Figure 6)^63^. Moreover, *P. aeruginosa* produces exoproteins such as PA-I lectin, which induce host barrier disruptions and have been clinically associated with intestinal colonization, particularly in immunosuppressed individuals, potentially explaining its sporadic presence in the gut^64^. In contrast, *B. cenocepacia* was mainly found in the gut. However, *B. cenocepacia* has primarily been studied as a respiratory pathogen in humans, and, to our knowledge, the gut has received little attention as a potential site of infection for this pathogen.

To determine differences in bacterial virulence in single and coinfection, virulence gene expression was monitored and, in *P. aeruginosa,* virulence-related genes were generally upregulated during coinfection (Figures 5 and 6). It is well-known that opportunistic bacteria such as *P. aeruginosa* can easily adapt to stress and very diverse environments. Coexistence with other microbes reduces nutrient availability at the infection site, resulting in both outcompeting and cooperative behaviors^65^. This could account for the change in gene expression observed in the global transcriptomic analysis, in which metabolic and biosynthetic pathways showed the greatest differential expression (Figure 6). Regarding virulence genes, the siderophore-encoding families pyoverdine- and pyochelin-encoding, which have been directly linked to *P. aeruginosa* virulence and have already been described in *B. cenocepacia-P. aeruginosa* interactions^66^ were upregulated during coinfection (Figure 6 and Table S6). Other upregulated genes in coinfection, such as *sodM* encoding a superoxide dismutase and the *pfeS/pfeR* two-component system, were shown to play a relevant role in the *in vivo* infections of *Bombyx mori* and *Vibrio harveyi*, respectively (Table S6)^67; 68^. When comparing virulence-related gene expression between acute and chronic strains in coinfection (Figure 5), *exoS* and *toxA* were the most upregulated for *P. aeruginosa* PAET1 and *P. aeruginosa* PAO1, respectively. Exoenzyme S, with both GTPase-activating and ADP-ribosyltransferase activitie^25^, disrupts host cell signaling, the cytoskeleton, and immune responses, and also plays a role in infection dissemination. This T3SS protein modulates host cell function to avoid bacterial detection while promoting infection persistence^69–71^. On the other hand, Exotoxin A inhibits eukaryotic protein synthesis by ADP-ribosylating elongation factor 2, leading to host cell death. Thus, it is highly expressed during acute infections, as it contributes to tissue damage and host immune dysfunction^33^, potentially explaining the lower hemocyte concentration in *P. aeruginosa* PAO1 coinfection compared with single infection (Figure 2J).

Unlike *P. aeruginosa*, most of the *B. cenocepacia* differentially expressed genes appeared to be upregulated during single infection. Focusing on its pathogenesis, overexpressed phenylalanine metabolism *paa* genes were shown to be involved in the *in vivo* virulence of *the Caenorhabditis elegans* infection model^33^, as well as to participate in the regulation of quorum-sensing-dependent virulence factors^33^. In addition, the overexpressed carbon metabolism-related *pgl* gene has been implicated in *Enterococcus faecalis* to take part in commensal and pathogenic interactions with the lepidopteran *Manduca sexta*^43^. Interestingly, *trxB,* encoding a thioredoxin-disulfide reductase, was also found to be upregulated in single infection, playing an important role in pathogen intracellular survival and oxidative stress defense^72^, which could be linked to the observed intracellular nature of *B. cenocepacia* in hemocytes (Figure 2 D and H). This extensive shift in gene expression during single infection (Figure 6E) might also be due to the fact that *B. cenocepacia* single-infected larvae lose viability only after 48 h and samples were consequently extracted after 30 h, while coculture samples were extracted after 15 h of infection. This suggests that at least some of the virulence genes of *B. cenocepacia* are likely expressed later.

Although fewer genes were overexpressed in *B. cenocepacia* coinfection (Figure 6D-E), some of them play a central role in its virulence. In fact, *cheA* and *cheW* genes were the only ones characterized in the bacterial chemotaxis KEGG classification and have been linked to bacterial adhesion in *Treponema denticola*^73^, and are also responsible for the correct flagellar function in *P. aeruginosa*^74^. In addition, the T2SS-related genes *gspD* and *gspG* are required for the correct function of the T2SS^75^, while *cblA* has been described to play a central role in *B. cenocepacia* adhesion in infection contexts^18; 34^. Similarly, virulence genes such as *cepR* and *cblA* were upregulated when assessed via RT-qPCR during coinfection (Figure 5), potentially contributing to the rapid decline in viability in coinfected larvae observed in Figure 1. Overall, these findings suggest that coinfection enhances the virulence of both *P. aeruginosa* PAO1 and *B. cenocepacia*, although RNA-seq with *P. aeruginosa* PAET1 in single and coinfection would also be required to obtain a clearer picture of the differences between acute and chronic *P. aeruginosa* strains. Nevertheless, its virulence gene expression, quantified via RT-qPCR, showed upregulation of all studied genes during coinfection, consistent with the observed larval lethality.

The present multifaceted investigation into single infections and coinfections of *B. cenocepacia* and *P. aeruginosa* demonstrates that coinfection significantly worsens infection outcomes. This is largely due to immunosuppression by *B. cenocepacia* and the enhanced virulence of *P. aeruginosa*. Additionally, acute *P. aeruginosa* PAO1 infection leads to rapid mortality and a strong immune response, while *P. aeruginosa* PAET1 chronic strain progresses more subtly but displays greater antimicrobial resistance, posing a significant long-term risk to host survival. While the *G. mellonella* model provides valuable insights in infection dynamics, it has limitations, particularly in its inability to support biofilm formation, a key feature of chronic infections. To advance our understanding, future research should focus on more complex *in vivo* models, particularly those with respiratory systems that better mimic human physiology. Overall, this work expands our understanding of the complex interplay between host responses and microbial communities in polymicrobial infections, reinforcing the need for therapeutic strategies that account for the complexity of these infections.

## Acknowledgements

This work was supported by grants PID2024-158933OB-100 funded by MCIN/AEI/10.13039/501100011033 and “ERDF A way of making Europe”, the CERCA programme and AGAUR-Generalitat de Catalunya (European Regional Development Fund FEDER) (2021SGR01545) and Catalan Cystic Fibrosis association. E.T. is a researcher of the ICREA Academia 2025 program. J.A-A. acknowledges MCIN for its financial support through a PRE2021-098703 grant funded by MCIN/AEI/10.13039/501100011033 and by the ESF+ “investing in your future”.

## Declaration of generative AI use

During the preparation of this manuscript, the authors used generative AI to improve the readability and language. Following the use of this tool, the authors reviewed and edited the content as needed and take full responsibility for the content of the published article.

## Author contributions statement

The manuscript was written through the contributions of all authors. Júlia Alcàcer-Almansa and Joana Admella performed all the experiments and wrote the manuscript. Núria Blanco-Cabra supervised the research, revised the experimental data and wrote the manuscript. Eduard Torrents directed and supervised the research, revised the experimental data, administered the project, acquired the funding and wrote the manuscript. All authors have approved the final version of the manuscript.

## Data availability statement

The data represented in all the figures of this study is available under the repository name “*Burkholderia cenocepacia* and *Pseudomonas aeruginosa* coinfection worsens infection outcomes by altering infection dynamics and compromising host immune effectiveness” at https://doi.org/10.34810/DATA2308. Raw sequencing data was deposited in the European Nucleotide Archive with the project accession code PRJEB120795.

