## Supplementary Information for "*Burkholderia cenocepacia* and *Pseudomonas aeruginosa* coinfection increases host mortality by altering infection dynamics and compromising host immune effectiveness"

**Table S1**. List of strains and plasmids used in this work.

| **Name** | **Description** | **Source** |
| --- | --- | --- |
| **Burkholderia cenocepacia** |  |  |
| J2315 | Wild type | ATCC BAA-245 |
| **Pseudomonas aeruginosa** |  |  |
| PAO1 | Wild type | ATCC 15692 |
| PA54 | Wild type, acute infection clinical strain | ^1^ |
| PA166 | Wild type, acute infection clinical strain | ^1^ |
| PAET1 | Wild type, clinical strain obtained from a patient with chronic CF infection. | ^1^ |
| PAET2 | Wild type, clinical strain obtained from a patient with chronic CF infection. | ^1^ |
| PAET4 | Wild type, clinical strain obtained from a patient with chronic CF infection. | ^1^ |
| MK171 | PAO1, chromosomically expressing eGFPmut3 | ^2^ |
| **Plasmids** | | |
| pETS134 | pETS130 carrying the *nrdA* promoter of *P. aeruginosa* | ^3^ |
| pETS248 | pETS130 with tetracycline resistance gene (Tc) carrying the *nrdB* promoter from *B. cenocepacia* and a blue fluorescent protein (BFP) | ^4^ |

**Table S2**. List of primers used in RT-qPCR experiments.

| **Gene** | **Protein** | **Forward Primer (5’-3’)** | **Reverse Primer (5’-3’)** |
| --- | --- | --- | --- |
| ***G. mellonella*** | | | |
| ***rel*** | Relish | TCCAAAAAGCACCCTACAATCG | GCACTTCGTAGCTCACATCTC |
| ***glo*** | Gloverin | AGATGCACGGTCCTACAG | GATCGTAGGTGCCTTGTG |
| ***mor*** | Moricin | GCTGTACTCGCTGCACTGAT | TGGCGATCATTGCCCTCTTT |
| ***nos*** | Nitic oxide synthase | ATGAAGGTGCTGAAGTCACAA | GCCATTTTACAATCGCCACAA |
| ***lysozyme*** | Lysozyme | TCCCAACTCTTGACCGACGA | AGTGGTTGCGCCATCCATAC |
| ***hemolin*** | Hemolin | CCCGAAGACGCTGGTGAATA | CGCACGTTCATTTGCTGTTC |
| ***18S*** | 18S ribosomal RNA | ATGGTTGCAAAGCTGAAACT | TCCCGTGTTGAGTCAAATTA |
| ***P. aeruginosa*** | | | |
| ***exoS*** | ExoenzymeS | GGCGGATGCGGAAAAGTAC | CTGACGCAGAGCGCGATT |
| ***toxA*** | ExotoxinA | ACATCAAGGTGTTCATCC | GACGAAGAAGGTGGCATC |
| ***vgrG2b*** | Type VI secretion system spike protein VgrG2b | CGCCAAGGTCGATATGAAACACAC | GTTCGGCGCCATCACGT |
| ***lasB*** | Elastase B | TGTCCAAACTCCCCAGCAAG | CGGATCGCTTTCAGTTCGTC |
| ***plcH*** | Hemolytic phospholipase C | TGAACTCGAACCGAACCAGG | GTTGGCTTCCGTATTGCTGG |
| ***gap*** | Glyceraldehyde-3-phosphate dehydrogenase | GAGGTTCTGGTCGTTGGTGT | GAGTGCACGGGGCTCTTC |
| ***B. cenocepacia*** | | | |
| ***cblA*** | Giant cable pilus | CTCTGATGTCGATGTCGGCT | CTACAGCTGCCTGAAGACCC |
| ***cepR*** | N-acylhomoserine lactone dependent regulatory protein CepR | GGCGCAGAACTACATCGAGA | CACCGACAATCCGAAATCGC |
| ***gspE*** | Type II secretion system ATPase GspE | TCGAGACGTTCTCGCTGAAG | GTTCTTCGTCAGGCTCCAGG |
| ***tssD*** | Type VI secretion system protein TssD | CGTCGTACCTGACCGACATC | GAGGTCGATGAACACCTCGG |
| ***pvdA*** | L-ornithine 5-monooxygenase | ACCGCTATCCGACGATTACC | CGAACAGGTCGAGTTCCCAC |
| ***rnd9*** | Resistance-nodulation-cell division efflux pump 9 | GGCAAGGAAGACGATTTCGC | GTTTGCGAAGAACACCGTCG |
| ***gyrB*** | DNA gyrase subunit B | CTCTGATGTCGATGTCGGCT | CTACAGCTGCCTGAAGACCC |

**Table S3. Health index score evaluation criteria.** The scoring system is based on the one previously published^5^.

| Category | Description | Score |
| --- | --- | --- |
| Activity | No movement | 0 |
|  | Minimal movement when stimulated | 1 |
|  | Movement when stimulated | 2 |
|  | Movement without stimulation | 3 |
| Cocoon formation | No cocoon | 0 |
|  | Partial cocoon | 0.5 |
|  | Full cocoon | 1 |
| Melanization | Complete | 0 |
|  | Black spots on brown larvae | 1 |
|  | 3 black spots on beige larvae | 2 |
|  | <3 black spots on beige larvae | 3 |
|  | None | 4 |
| Survival | Dead | 0 |
|  | Alive | 2 |

**Table S4**. Bacterial quantification from the fat body, gut and hemolymph through plating and confocal microscopy images of larval dissections. The data shown corresponds to the mean of the bacterial counts for each group. Fat body and gut bacterial quantifications were performed through ImageJ analysis of three confocal images per group derived from larvae dissections. Hemolymph bacterial quantification was performed through hemolymph extraction and plating in TSA plates. The following abbreviations were used: *P. aeruginosa* PAO1 (PAO1), *P. aeruginosa* PAET1 (PAET1), *B. cenocepacia* (BC). Coinfection was noted as “Co-” before the corresponding strain. Each condition was tested with a minimum of three replicates.

|  | |  | | **Number of bacteria in the fat** | | **Number of bacteria in the gut** | | **CFU/mL in the hemolymph** | | |
| --- | --- | --- | --- | --- | --- | --- | --- | --- | --- | --- |
|  |  | | **Early infection** | | **Late infection** | **Early infection** | **Late infection** | | **Early infection** | **Late infection** |
| Single  infection | **BC** | | 271 | | 21 | 91 | 1373 | | 5x10^3^ | 4.8x10^4^ |
|  | **PAO1** | | 0 | | 600 | 0 | 227 | | 120 | 2.2x10^8^ |
|  | **PAET1** | | 752 | | 25 | 1487 | 15 | | 4.5x10^6^ | 9x10^7^ |
| CoPAO1 | **BC** | | 171 | | 3 | 155 | 23 | | 5.5x10^5^ | 1.4x10^5^ |
|  | **PAO1** | | 0 | | 363 | 0 | 501 | | 95 | 2.4x10^7^ |
| CoPAET1 | **BC** | | 142 | | 1 | 6 | 10 | | 3.9x10^5^ | 8.6x10^5^ |
|  | **PAET1** | | 97 | | 42 | 33 | 34 | | 100 | 1.1x10^6^ |

**Table S5.** Immunity-related *G. mellonella* genes differentially expressed in larvae after single infection with *P. aeruginosa*, *B. cenocepacia* or in coinfection.

| **Gene ID** | **Gene description** | **Log2 Fold change** |
| --- | --- | --- |
| **DEGs in larvae infected with *P. aeruginosa* PAO1 single infection** | | |
| LOC113515727 | Hemolectin | 4.6 |
| LOC113519878 | Proteoglycan-like sulfated glycoprotein papilin | 1.6 |
| LOC113509888 | Toll-like receptor 3 | 3.5 |
| LOC113517945 | Phenoloxidase subunit 1 (PPO1) | 3 |
| LOC113515116 | Phenoloxidase subunit 2 (PPO2) | 3.2 |
| LOC113516387 | Apolipophorin-3-like | 2.3 |
| LOC113522331 | Cathepsin O-like | 1.7 |
| LOC113521879 | A disintegrin and metalloproteinase with thrombospondin motifs 17-like | 3.1 |
| LOC113522639 | Hatching enzyme-like protein (hel) | 3.3 |
| LOC113522669 | Astacin-like metalloprotease toxin 5 | 4.9 |
| **DEGs in larvae infected with *P. aeruginosa* PAO1 coinfection** | | |
| LOC113522123 | Heat shock protein Hsp-12.2-like | 1.5 |
| LOC113523440 | Gallerimycin-like | 1.5 |
| LOC113510422 | Gloverin-like | 2.3 |
| LOC128201089 | Gloverin-like | 1.7 |
| LOC113515290 | Lysozyme | 1.6 |
| LOC128201155 | Heat shock protein 68-like | 1 |
| LOC113509608 | Moricin | 1.3 |
| **DEGs in larvae infected with *B. cenocepacia* single infection** | | |
| LOC113514843 | CD63 antigen-like | 1.5 |
| LOC113518019 | Gamma-interferon-inducible lysosomal thiol reductase-like | 2 |
| LOC113517748 | Apomucin-like | 1.8 |
| LOC113515559 | CLIP domain-containing serine protease B4 | 2.7 |
| **DEGs in larvae infected with *B. cenocepacia* coinfection** | | |
| LOC113513204 | Spatzle-1 (spz1) | 1.5 |
| LOC113518960 | Peroxidase-like | 1.7 |
| LOC113511064 | Oxidative stress-induced growth inhibitor 1-like | 1 |
| LOC113511903 | Catalase-like | 2.5 |
| LOC113516037 | Spaetzle domain-containing protein 3 (spz3) | 2.2 |
| LOC113511546 | Spaetzle 5 Toll-1 receptor (spz5) | 1.6 |
| LOC113515637 | Relish | 1.2 |

**Table S6.** Secretion system, siderophore production, quorum sensing, virulence and antibiotic resistance-related genes that were differentially expressed in *P. aeruginosa* PAO1 when coinfected with *B. cenocepacia* resulting from RNA-sequencing analysis. The references (Ref) include articles where these genes have been described in virulence contexts, and the “Species” category corresponds to the bacterial species in which the gene has been described to participate in virulence processes.

| **Gene name** | **Description** | **Species** | **Log2 Fold change** | **Ref** |
| --- | --- | --- | --- | --- |
| **Secretion systems** | | | | |
| ***pscF*** | **Type 3 secretion system needle filament protein** | *P. aeruginosa* | 1.27 | ^6^ |
| **Siderophores and hemophores** | | | | |
| ***pvdA*** | **L-ornithine N(5)-monooxygenase** | *P. aeruginosa* | 1.74 | ^7; 8^ |
| ***pvdD*** | Pyoverdine synthetase D | *P. aeruginosa* | 1.1 |  |
| ***pvdE*** | Pyoverdine biosynthesis protein PvdE | *P. aeruginosa* | 1.5 |  |
| ***pvdF*** | **Phosphoribosylglycinamide formyltransferase 1** | *P. aeruginosa* | 1.1 |  |
| ***pvdG*** | Pyoverdine biosynthesis thioesterase PvdG | *P. aeruginosa* | 1.2 |  |
| ***pvdH*** | L-2,4-diaminobutyrate:2-ketoglutarate 4-aminotransferase, PvdH | *P. aeruginosa* | 1.3 |  |
| ***pvdJ*** | Pyoverdine biosynthesis protein PvdJ | *P. aeruginosa* | 1.3 |  |
| ***pvdL*** | Non-ribosomal peptide synthetase PvdL | *P. aeruginosa* | 1.6 |  |
| ***pvdN*** | Pyoverdine biosynthesis protein PvdN | *P. aeruginosa* | 1.5 |  |
| ***pvdO*** | Pyoverdine biosynthesis-related protein PvdO | *P. aeruginosa* | 1.4 |  |
| ***pvdP*** | Pyoverdine biosynthesis-related protein PvdP | *P. aeruginosa* | 1.3 |  |
| ***pvdQ*** | Acyl-homoserine lactone acylase PvdQ | *P. aeruginosa* | 1.5 |  |
| ***pvdR*** | Pyoverdine export membrane fusion protein PvdR | *P. aeruginosa* | 1.1 |  |
| ***pvdS*** | Sigma factor PvdS | *P. aeruginosa* | 1.1 |  |
| ***pvdT*** | Pyoverdine export ATP-binding/permease protein PvdT | *P. aeruginosa* | 1 |  |
| ***pchE*** | Pyochelin synthetase PchE | *P. aeruginosa* | 1.1 | ^9^ |
| ***pchF*** | Pyochelin synthetase PchF | *P. aeruginosa* | 1 |  |
| ***pchR*** | Regulatory protein PchR | *P. aeruginosa* | 1.5 |  |
| ***tonB2*** | Protein tonB2 | *P. aeruginosa* | 5 | ^10^ |
| ***fptA*** | **Fe(3+)-pyochelin receptor** | *P. aeruginosa* | 1.5 | ^11^ |
| ***fpvA*** | Ferripyoverdine receptor | *P. aeruginosa* | 1.7 | ^12^ |
| ***fpvB*** | Second ferric pyoverdine receptor FpvB | *P. aeruginosa* | 1.4 | ^13^ |
| ***pfeA*** | Ferric enterobactin receptor | *P. aeruginosa* | 1.7 | ^14^ |
| ***pirA*** | Ferric enterobactin receptor PirA | *P. aeruginosa* | 1.3 | ^15^ |
| ***hasAp*** | Heme acquisition protein HasAp | *P. aeruginosa* | 2.4 | ^16^ |
| ***hasE*** | 7-dimethylallyltryptophan synthase HasE | Serratia marcescens | 2.4 | ^17^ |
| ***hasR*** | Heme uptake outer membrane receptor HasR | *P. aeruginosa* | 2.1 | ^18^ |
| ***phuR*** | Heme/hemoglobin uptake outer membrane receptor PhuR | *P. aeruginosa* | 1.6 | ^19^ |
| ***phuT*** | Heme-transporter PhuT | *P. aeruginosa* | 1.39 | ^20^ |
| **Quorum sensing** | | | | |
| ***lasI*** | Acyl-homoserine-lactone synthase | *P. aeruginosa* | 1.6 | ^21^ |
| ***rhlB*** | ATP-dependent RNA helicase RhlB | *P. aeruginosa* | 2.8 | ^22^ |
| ***rhlI*** | **Acyl-homoserine-lactone synthase** | *P. aeruginosa* | 1.52 | ^23^ |
| ***ambA*** | LysE-type transporter | *P. aeruginosa* | 2.37 | ^24^ |
| ***ambB*** | non-ribosomal peptide synthetase AmbB | *P. aeruginosa* | 1.8 |  |
| **Virulence** | | | | |
| ***serA*** | **D-3-phosphoglycerate dehydrogenase** | *P. aeruginosa* | 1 | ^25^ |
| ***trpA*** | Tryptophan synthase alpha chain, TrpA | *Pseudomonas cannabina* in plants | 1.2 | ^26^ |
| ***sodM*** | **Superoxide dismutase** | *P. cannabina* in *Bombyx mori* | 1.7 | ^27^ |
| ***aprA*** | Alkaline protease | *P. aeruginosa* | 1.3 | ^28^ |
| ***aprE*** | Alkaline protease secretion protein AprE | *P. aeruginosa* | 1.7 | ^29^ |
| ***aprF*** | Alkaline protease secretion protein AprF | *P. aeruginosa* | 1.9 |  |
| ***pfeR*** | Transcriptional activator protein PfeR | *Vibrio harveyi* in fish blood | 1.3 | ^30^ |
| ***pfeS*** | Sensor protein PfeS |  | 1.4 |  |
| ***foxA*** | Metal-pseudopaline receptor CntO | *P. aeruginosa* | 1.6 | ^31^ |
| ***pasP*** | Small protease PasP | *P. aeruginosa* | 1.3 | ^32^ |
| ***ptxS*** | HTH-type transcriptional regulator PtxS | *P. aeruginosa* | 1.1 | ^33^ |
| ***fumC1*** | Fumarate hydratase class II 1 | *P. aeruginosa* | 1.75 | ^34^ |

**Figure S1. Bacterial dose optimization.** Kaplan-Meier survival curves of *G. mellonella* infected with different doses of **A)** *B. cenocepacia* (BC), **B)** *P. aeruginosa* PAET1 as the representative chronic strain (PAET1) and **C)** Coinfection of BC and PAET1. *P. aeruginosa* PAO1 as the representative acute strain dose optimization has been previously published by our lab ^35^. Doses are expressed in CFU/larva. Statistical significance was established through a log-rank (Mantel-Cox) survival test comparing the control group with infected groups.

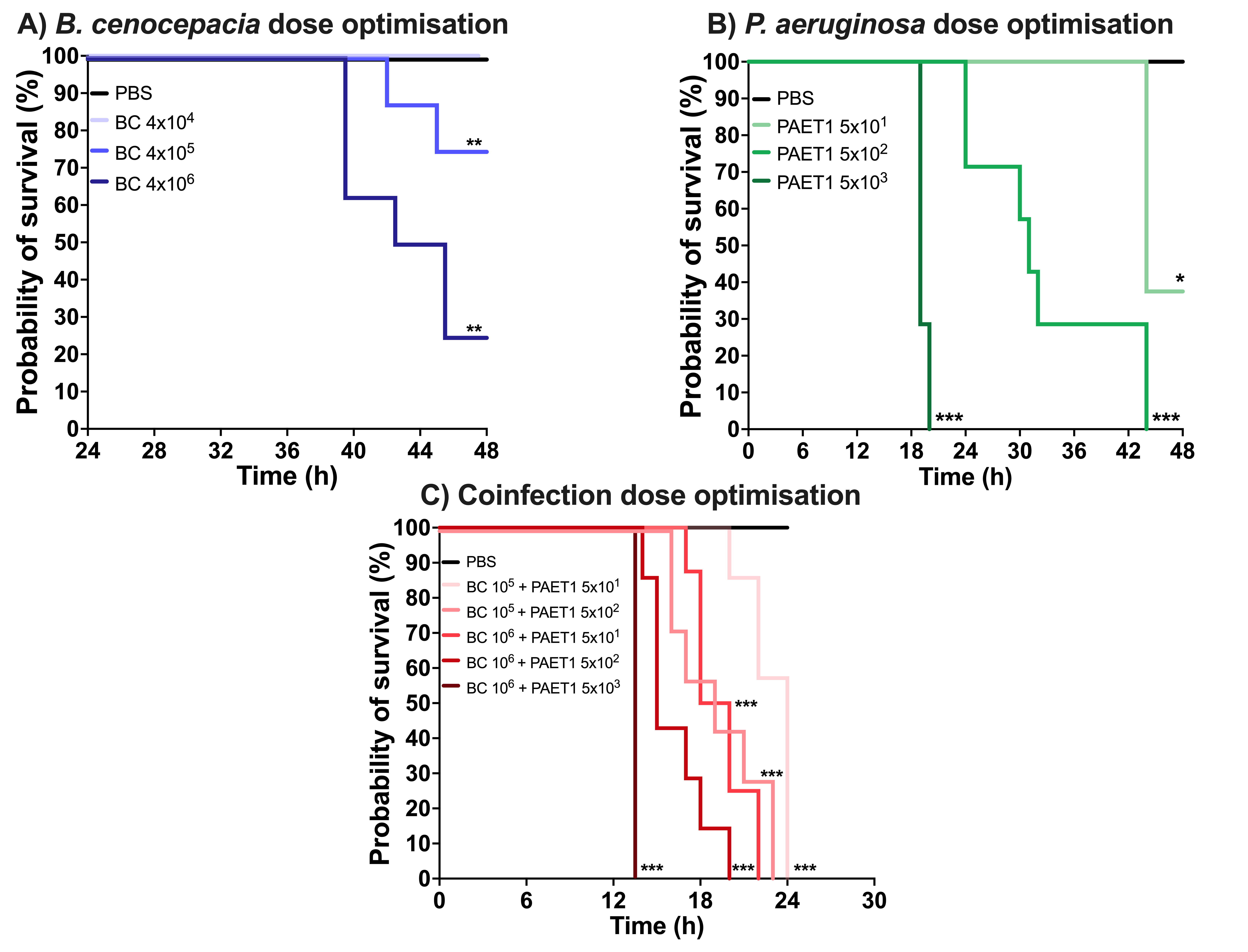

**Figure S2. A) Larval health according to the health index score B) Bacterial quantification in whole larvae.** Data is presented as the mean ± SD for each condition. The following abbreviations were used: *P. aeruginosa* PAO1 (PAO1), *P. aeruginosa* PAET1 (PAET1), *B. cenocepacia* (BC). Coinfection was noted as “Co-” before the corresponding strain. For early and late infection timepoints, see materials and methods. Statistical significance was established via two-way ANOVA. The statistical significances that appear in the graph correspond to each group at early *vs.* late infection stages (dotted line), and for a specific stage, each group against control (full line) or individual groups *vs.* coinfection (dashed line), where **p*<0.05, ***p*<0.01, ****p*<0.001 and *****p*<0.0001. $ corresponds to significant differences (****) in all comparisons between the late infection stage of control (PBS) *vs.* the rest of groups.

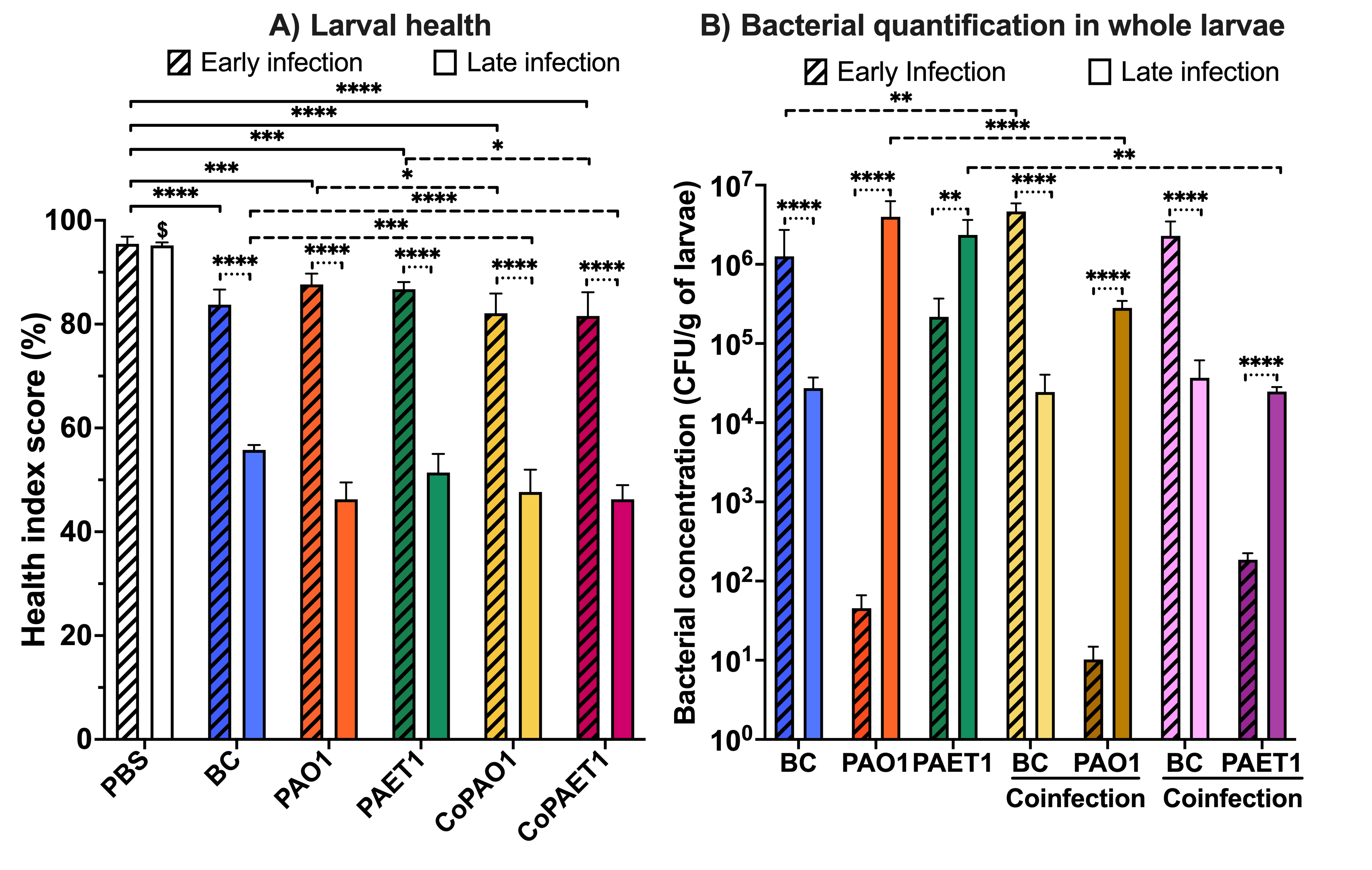

**Figure S3.** Confocal images of the hemolymph from larvae injected with PBS as a control group at early **(A)** and late **(B)** infection stages. **C)** Image of the whole *G. mellonella* body when infected with PBS. Hemocytes can be seen in red as they were stained with FM^TM^ 4-64 (Invitrogen). Scale bar corresponds to 20 μm.

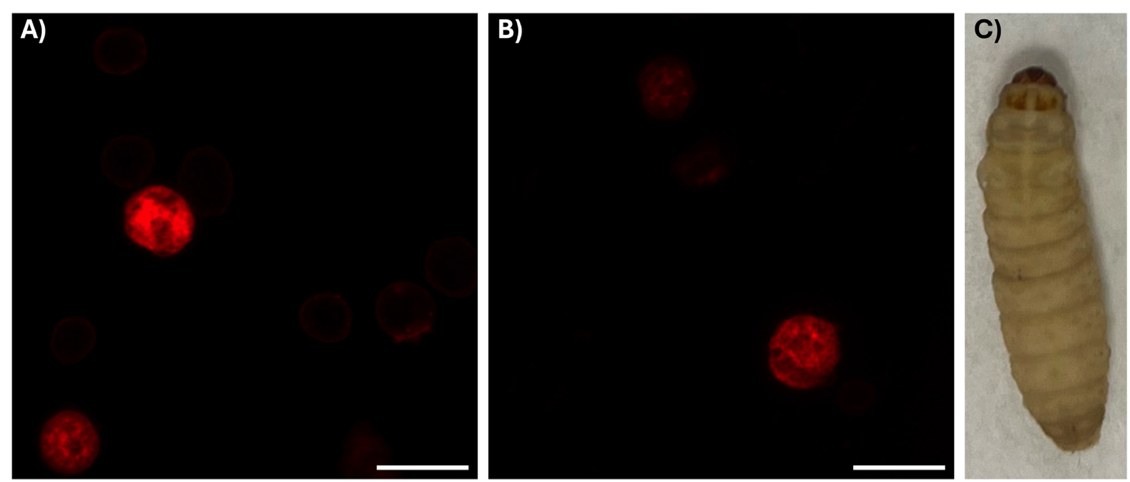
